# Automated Knowledge Graph Construction for CAR T Cell Receptor Design via Hybrid Text Mining

**DOI:** 10.64898/2026.04.06.716719

**Authors:** Haomiao Luo, Difei Tang, Anya Zivanov, Natasa Miskov-Zivanov

## Abstract

Designing next-generation Chimeric Antigen Receptors (CARs) requires a systematic understanding of intracellular signaling domains and their downstream biological effects, yet no comprehensive knowledge resource currently exists for this purpose. Here, we present an automated workflow that integrates multiple natural language processing and large language model tools to extract biomolecular interactions from PubMed literature and assemble them into a CAR T cell signaling knowledge graph. Our pipeline combines REACH, INDRA, and Llama 3 across 15 targeted search queries, yielding a directed multi-relational graph of ∼7,500 unique interactions among ∼1,800 entities, including proteins, biological processes, and chemicals. We further demonstrate that queries incorporating biological process ontology terms retrieve more interaction-rich papers than protein-name-only searches, offering practical guidance for future literature mining efforts. The resulting knowledge base provides a structured foundation for predicting T cell phenotypes and prioritizing intracellular domain candidates for CAR design, with broader applicability to knowledge-driven inference in immunotherapy research.

## 1 INTRODUCTION

Chimeric Antigen Receptor (CAR) T cell therapy has emerged as a promising immune therapy for hematology malignancy [1]. CAR consists of three major components: a single chain variable fragment (scFv) responsible for antigen recognition, a transmembrane domain, and intracellular domains (ICDs) that govern downstream signaling. The ICDs contain signaling motifs that modulate the cellular response such as T cell activation, proliferation, and persistence. Although achieving desired functions with receptor engineering, existing FDA-approved CAR T cell therapies are associated with significant adverse effects, such as cytokine release syndrome (CRS) and immune effector cell-associated neurotoxicity syndrome (ICANS) [2].

To overcome this drawback and discover next-generation CARs, recent efforts have focused on creating a large library of candidate receptors. Engineering the third generation CARs is more complex than earlier generations since it requires a combination of more co-stimulatory domains. T cells with different codomains exhibit variations in expansion, lifespans, and cytotoxicity. The positions of codomains further influences the fate of these cells. To navigate this complexity, high-throughput approaches often referred to as “CAR pooling” were proposed, which enable the screening of hundreds of thousands of domain combinations and sequences [3, 4]. Those methods typically involve high throughput experiments providing meaningful data resource for post-analysis.

While CAR pooling facilitates the experimental identification of promising candidates, it does not fully address the challenge of prioritizing candidate signaling motifs from the vast space of possible intracellular domain configurations. Recent study employed machine learning to decode CAR T cell phenotype and predict the cytotoxicity and stemness of CAR T cells [5]. However, these approaches typically rely on supervised learning frameworks that require extensive labeled datasets, which are often limited in size and scope. Consequently, such models may struggle to generalize beyond the predefined domain combinations represented in the training data.

To address these limitations, we propose constructing a comprehensive knowledge graph that connects receptors with resulting biological processes through signaling pathways. We developed a customized text-mining workflow to extract intracellular interactions from literature available in PubMed Central. Specifically, we used targeted search queries that include names of ICD candidates, biological processes, and their molecular markers. Next, we utilized a hybrid approach to improve accuracy of extracted interactions from retrieved texts. Finally, we evaluated advantages and drawbacks of database-driven filtering for removing low-confidence interactions.

## 2 METHODOLOGY

The steps of our workflow are illustrated in Figure 1 and detailed in the following, from creating search queries, retrieving relevant literature, using natural language processing (NLP) and large language model (LLM) tools to extract interactions from the literature, to generating and analyzing the network of all extracted interactions.

**Figure 1.**
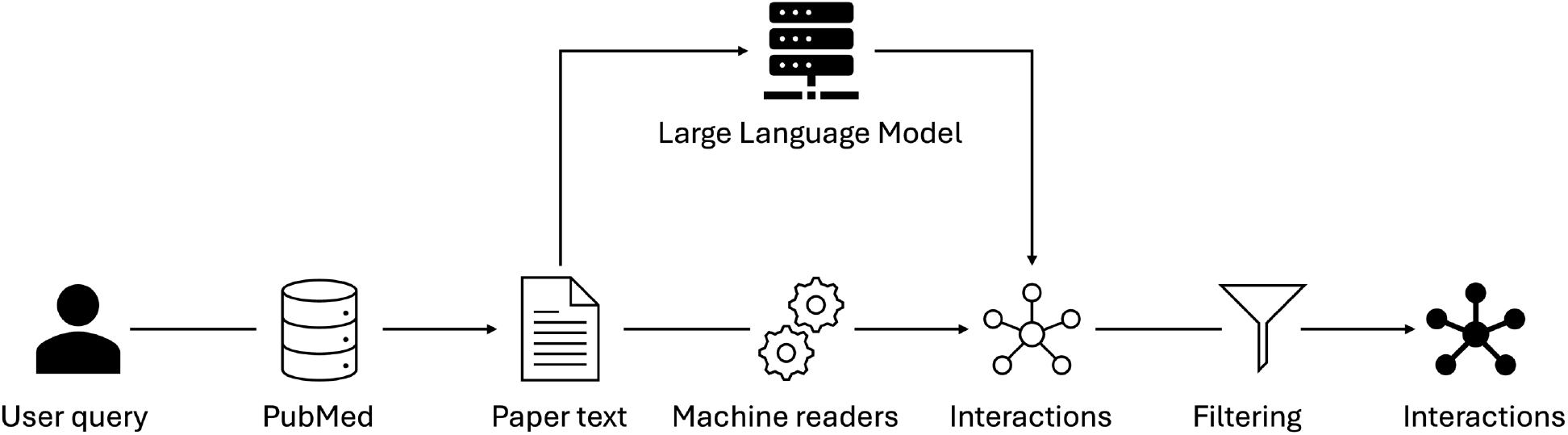
Pipeline for automatically extracting interaction from published literature for the biomolecular interaction knowledge base.

### 2.1 Query design

PubMed relies on combining query-document relevancy score, publication date, and type with the-state-of-the-art machine learning model to find the best matched documents to the query [6, 7]. Therefore, by varying search queries in PubMed we can obtain different document corpora, which in turn could influence the biological event extraction from biomedical literature [8, 9].

We created 15 different search queries using the following procedure (Figure 2). For the 40 ICD candidates from across the human proteome [4], we included additional common synonyms, resulting in 118 different terms in the \ domain candidates (DCs) group. The second and third group of terms were created from the list of relevant biological processes (BPs), and their corresponding process markers (PMs). To form queries, all terms within each group, DC, PM, and BP, were connected using “or” operator, which means that for a paper to be selected, at least one of the terms from the same group should be found in the paper (e.g., one of the included synonyms for at least one of the DC candidates). We then considered five different logical expressions, either the “or’-ed expression from a single group, or two or all three groups connected with an “and” operator: DCs only, DCs and PMs, DCs and PMs and BPs, PMs only, and PMs and BPs. Next, as context is important for mechanistic models [10], each of these five combinations was considered for three different contexts: CAR T cell, T cell, and no context, thus resulting in total 15 different search queries.

**Figure 2.**
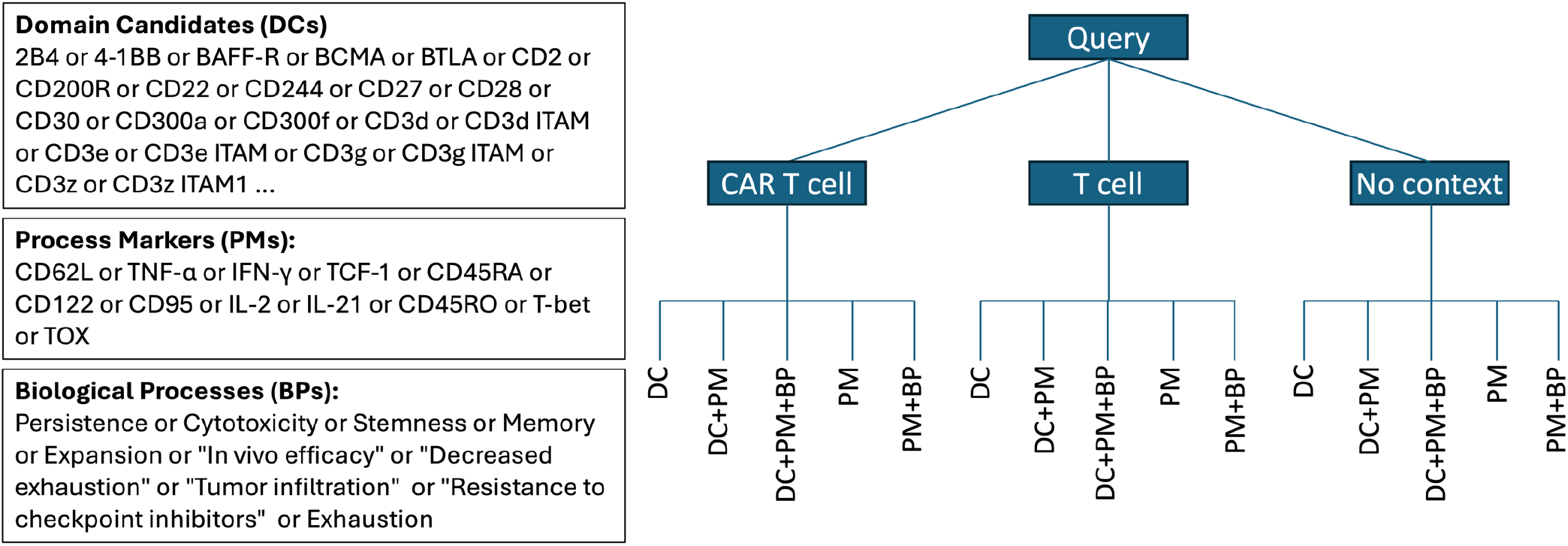
All terms used for creating queries(left) and an outline of the process for creating final 15 queries (right). Terms within the same group are combined with operator “or” and different groups are connected with operator “and”. DCs are the candidates considered for intracellular receptor domains. Process markers (PMs) are biomarkers of the biological process (BPs) of interest for CAR T cells.

### 2.2 Document retrieval

We used PubMed E-utilities API to retrieve the relevant Paper IDs where the sorting method is set to “best match” and database is configured with PubMed Central (PMC). The max number of retrieved papers is set to 2,000. Both the INDRA and E-utilities APIs are used to collect the full texts for retrieved PMC IDs. The acquired PMC IDs are used to fetch the corresponding XML files from PMC, compatible with REACH input/output requirements, while the text files accessed via INDRA API are further used for Llama 3 text generation.

### 2.3 Interaction extraction

After retrieving paper corpora for all search queries, we utilized the REACH tool [11] and INDRA database [12, 13] to extract interactions from every paper. Additionally, we used INDRA and Llama 3 [14] to extract interaction from those papers where REACH was unable to find any interactions. We used the INDRA database to retrieve text files of the documents that are not readable by REACH. The Llama 3 text-generation model is used to extract the interactions through the Hugging Face platform. Specifically, we used in-context learning to extract interactions. Enhanced by the inductive head of the language model, the few-shot learning prompt (Figure 3) is used to improve the inference of Llama 3. The learning examples in the prompt include the input unstructured text and a structured BioRECIPE format [15] with the information about biological regulator entity, biological regulated entity, an interaction between them, and additional attributes that further specify the regulator, regulated and the interaction. A parser is also developed to fetch the text information to BioRECIPE interaction spreadsheet. Since text generated from Llama 3 is based on the probability of the tokens and might have “hallucinations”, we also created scripts to post-process the generated text.

**Figure 3.**
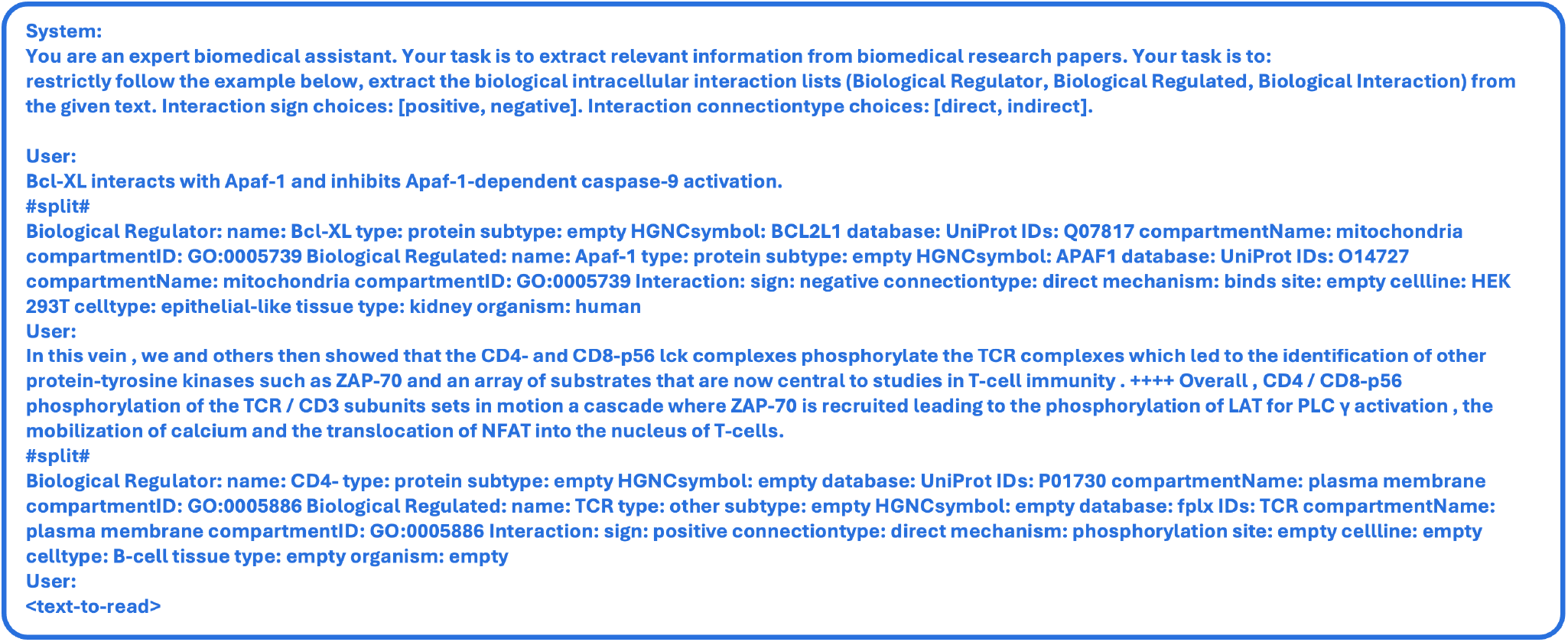
The prompt for the few-shot learning approach used for Llama 3.

### 2.4 Interaction filtering

While the existing event extraction tools are useful, recent reports of their precision and recall highlight that further improvement are needed and they may not be fully reliable [13, 16]. Therefore, to keep only the high-confidence interactions, we also created a version of each corpus using a filtering tool FLUTE [17] that scores the reliability of the interactions based on the interaction databases such as STITCH [18], STRING [19], and gene ontology (GO) [20] and identifies the most reliable interactions based on the set threshold.

### 2.5 Event graph construction and evaluation

To further evaluate the information that we retrieved through our literature search, we created a graph from all extracted interactions, using the Python package NetworkX [21]. This enables analysis of the relationships and signaling pathways between co-signaling domains and biological processes. We used the centrality metric to measure how many nodes the codomains connect with and we used the PageRank score to assess how important each codomain is in this network. Additionally, each node (protein, gene, biological process, etc.) in our knowledge graph was converted into a numerical vector using the Node2Vec algorithm, which learns representations by simulating random walks through the graph. Nodes that are more similarly connected in the graph have similar vectors. Next, we used principal component analysis (PCA) to reduce those higher dimensional vectors down to two dimensions so they can be plotted, with similar nodes appearing closer together.

## 3 RESULTS

### 3.1 Influence of the context in queries on corpus relevance

Each query returned a list of PubMed Central IDs of relevant papers, and the number of PubMed hits varied from ∼300 for the query that included “CAR T cell” context and DC+PM terms to almost 2M papers for the query without context and PM+BP terms (Figure 4(a)). As expected, our results confirmed that queries without any context return the largest number of papers, followed by queries with “T cell” context. Since CAR T cell research is more recent, queries with “CAR T cell” context led to the smallest number of papers.

**Figure 4.**
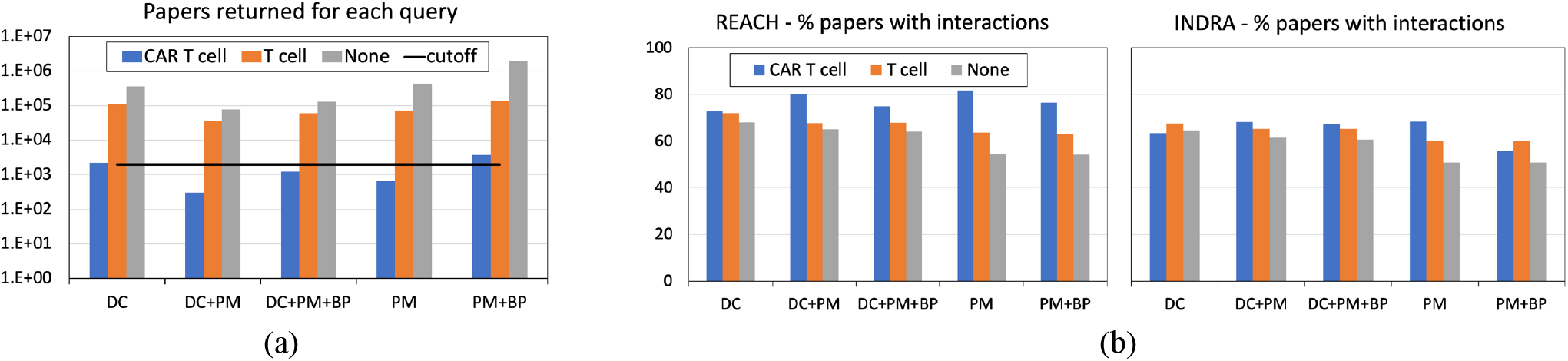
Papers retrieved with the 15 search queries, for the three contexts, CAR T cell, T cell, and None, and the five combinations of terms, CD only, CD+PM, CD+PM+BP, PM only, PM+BP. a) Total papers retrieved from PubMed and the cutoff for machine reader processing (line); b) % papers from the top 2000 retrieved from PubMed in which REACH (left) and INDRA (right) found at least one interaction.

As some queries resulted in 10^5^-10^6^ hits in PubMed, we limited the maximum number of papers to be processed by readers to 2,000 (Figure 4(a), line). For each query, using PubMed’s sort by relevance feature, we selected first 2000 (where available) paper IDs. Following this procedure, the total number of unique papers found across all queries was 10,060.

Next, for each paper corpus resulting from individual query, we evaluated the number of papers in which REACH and INDRA found interactions (Figure 4 (b)). When compared to total PubMed hits, using only the top hits led to almost opposite ordering in terms of the number of papers from which interactions were extracted, more CAR T cell papers resulted in extracted interactions. A possible explanation for this is that a more targeted search leads to papers with more signaling details.

We also investigated the number of extracted interactions from papers in each corpus jointly for REACH and INDRA (Table 1). The T cell context appears to lead to the largest number of extracted interactions, while the effect of the term group (DCs or PMs or BPs) on the amount of found interactions is less clear and varies across used contexts. As expected, the largest number of interactions with DCs is always found for the query that includes DCs only, since it explicitly searches for DCs, while being less restrictive than queries that also include other terms. Still most DC synonyms are found when no context is included in queries.

**Table 1.** Summary of found interactions for each query.

|  | QUERY | #Total interactions | #Interactions with DCs | #DC synonyms found in interactions |
| --- | --- | --- | --- | --- |
| CAR T cell | DC | 1880 | 639 | 44 |
|  | DC+PM | 418 | 130 | 17 |
|  | DC+PM+BP | 1353 | 470 | 39 |
|  | PM | 890 | 202 | 23 |
|  | PM+BP | 2381 | 581 | 40 |
| T cell | DC | **5662** | **2984** | 47 |
|  | DC+PM | **5172** | 2177 | 42 |
|  | DC+PM+BP | **5180** | 2368 | 46 |
|  | PM | 3787 | 831 | 32 |
|  | PM+BP | 3854 | 880 | 34 |
| None | DC | **5116** | **2675** | **61** |
|  | DC+PM | **5022** | 2009 | **59** |
|  | DC+PM+BP | 4880 | 2046 | **55** |
|  | PM | 2431 | 384 | 29 |
|  | PM+BP | 2466 | 401 | 30 |

To use the collected information in making predictions or explaining the behavior of CAR T cells, it is important to also account for the context within which the extracted interactions were mentioned in the papers. Therefore, we summarized the contextual information for extracted interaction list (Table 2). These results show that queries with “T cell” term included result in more consistent context information when compared to the other two context terms we used. Queries with “CAR T cell” lead to interactions that belong to more diverse cell types. When no context term is included in the query, the collected interactions are scattered across many different cell types.

**Table 2.** The top 10 cell types most often found in interactions extracted from the retrieved paper corpora and the number of interactions that include them for each corpus.

| CONTEXT FOUND IN INTERACTIONS \ QUERY TERMS | CAR T cell |  |  |  |  | T cell |  |  |  |  | No context |  |  |  |  |
| --- | --- | --- | --- | --- | --- | --- | --- | --- | --- | --- | --- | --- | --- | --- | --- |
|  | DC | DC+PM | DC+PM+BP | PM | PM+BP | DC | DC+PM | DC+PM+BP | PM | PM+BP | DC | DC+PM | DC+PM+BP | PM | PM+BP |
| T cell | 702 | 151 | 509 | 316 | 1046 | 1703 | 1592 | 1659 | 1171 | 1216 | 1238 | 1217 | 1242 | 495 | 496 |
| regulatory T cell | - | 25 | - | - | - | 479 | 363 | 410 | 218 | 225 | 400 | 324 | 321 | - | - |
| CD8-Positive T-Lymphocytes | 127 | 30 | 105 | 65 | 193 | 444 | 364 | 447 | 284 | 319 | 337 | 275 | 293 | - | - |
| CD4-Positive T-Lymphocytes | - | - | - | - | - | 581 | 743 | 685 | 421 | 448 | 395 | 565 | 539 | 198 | 203 |
| lymphocyte | - | - | - | - | - | - | - | - | - | - | - | - | - | 134 | 135 |
| B cell | 139 | - | 87 | - | 152 | 342 | 270 | 292 | 215 | 220 | 431 | 346 | 325 | - | 104 |
| natural killer cell | 118 | 33 | 89 | 64 | 228 | - | - | - | - | - | - | - | - | - | - |
| neoplastic cell | 208 | 41 | 153 | 89 | 361 | - | - | - | - | - | - | - | - | - | - |
| leukocyte | - | - | - | 49 | - | - | - | - | - | - | - | - | - | 154 | 149 |
| neutrophil | - | - | - | - | - | - | - | - | - | - | - | - | - | 106 | - |

### 3.2 Evaluation of the use of LLMs in the workflow

We used Llama 3 to generate events from those papers where REACH and INDRA did not find any interactions (849 unique papers). LLMs have superior performance on comprehending the literature and importantly, can also identify in texts the rich contextual information such as cell line, cell type, and organism. However, the raw output of LLMs is inherently unstructured, and therefore we used the prompt to instruct Llama 3 to output structured text.

### 3.3 Creation and analysis of CAR T cell knowledge graph

The knowledge graph was created by combining interaction lists obtained from REACH, INDRA, and Llama 3 across all 15 queries (Figure 5). The aggregated result is a directed multi-relational graph, with 7501 unique interactions and 1841 unique entities, mainly proteins, biological processes, and chemicals. In the graph, the protein-protein interactions are the dominant interaction type, whereas protein-chemical interactions are notably sparse. There is also a significant variability in the connectivity of different DCs within the graph (Table 3).

**Table 3.**
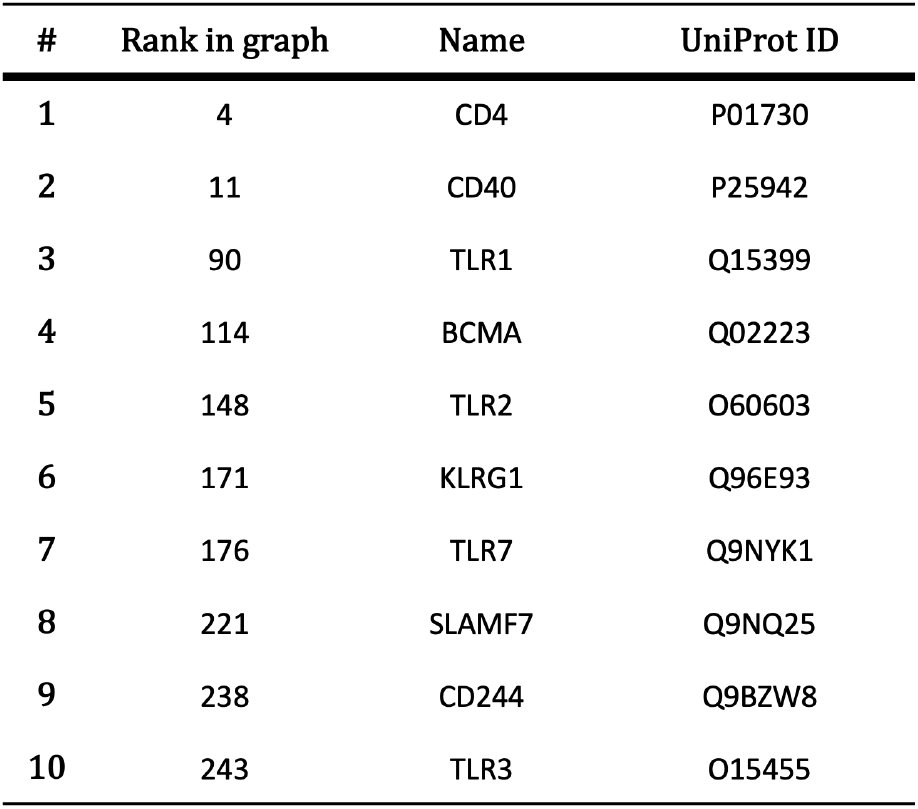
Top 10 DCs found in the generated knowledge graph.

**Figure 5.**
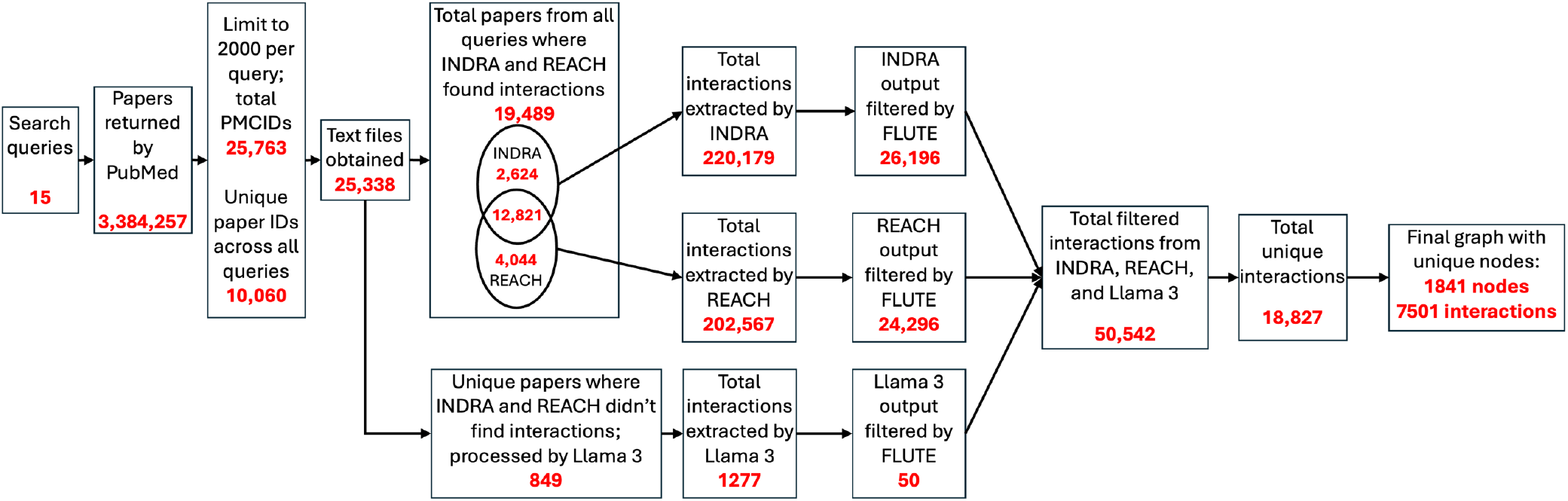
The workflow of collecting interactions for the knowledge graph from all 15 paper corpora, by combining outputs from INDRA, REACH and Llama 3, and filtering with FLUTE, with the number of interactions in each step.

Figure 6 shows the PCA projection of node embeddings into two dimensions, where the distance between points in the plot indicates similar connectivity in the knowledge graphs, that is, which DCs are “central” to the signaling network versus which ones occupy more peripheral or unique roles. As can be seen in the figure, the majority of DCs cluster in the top-right dense region, suggesting that most of them share similar interaction neighborhoods, connecting to many of the same proteins and biological processes. CD28 and SYK appear as outliers (far left cluster), indicating that they have very distinct connectivity patterns from the rest of the network. Similarly, CD27, KIR3DL2/3, and LAG3 are outliers in the bottom right, while CXADR and CD244 are also isolated as visible in the zoomed view, which again may indicate sparse or atypical connections in the graph.

**Figure 6.**
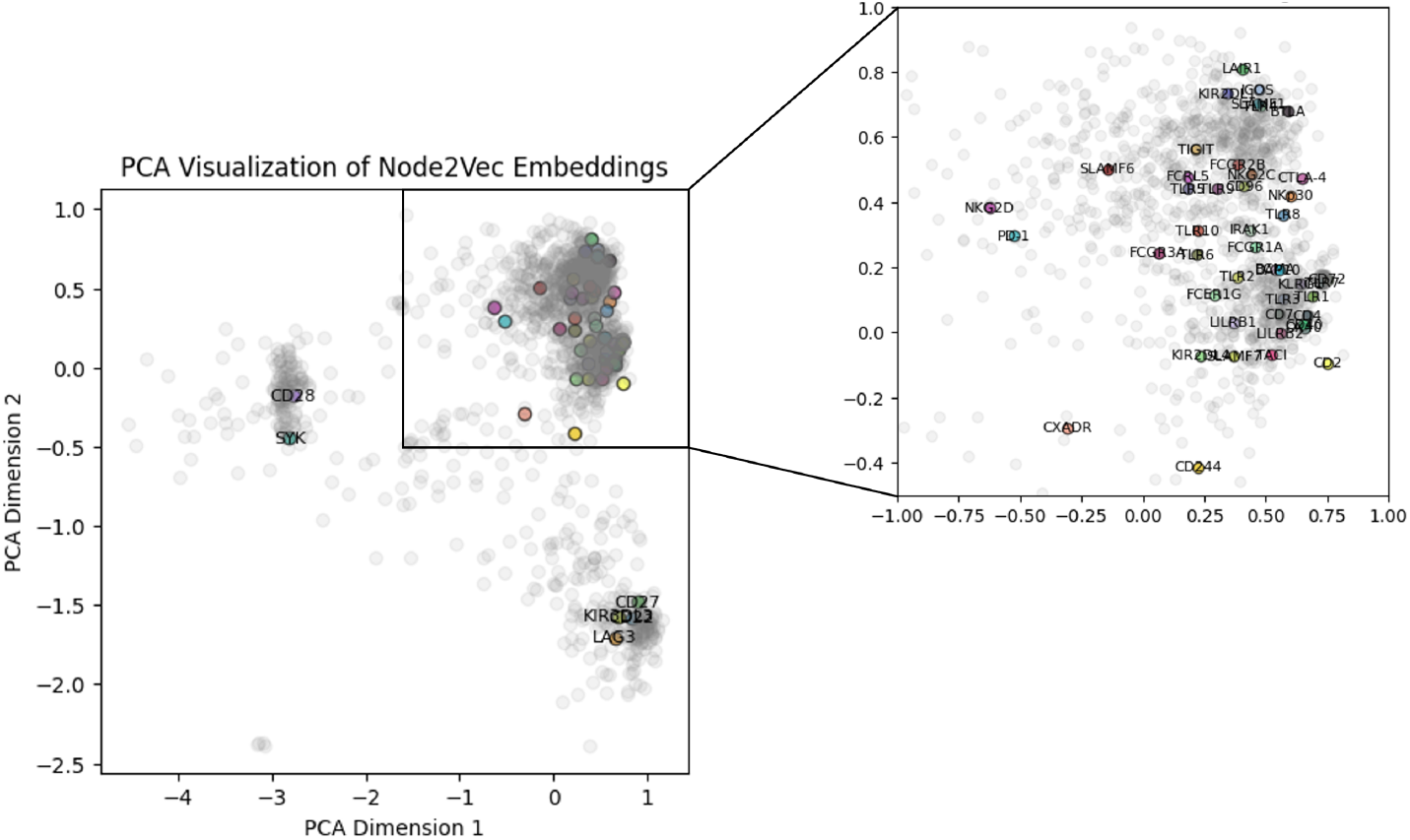
Principal Component Analysis of the Node2Vec embedding of the created CAR T cell literature-based knowledge graph. The grey dots are all graph nodes, while the colored and labeled nodes are DCs.

## 4 CONCLUSION

In this work, we integrated existing tools into a novel workflow to construct a knowledge graph of signaling in CAR T cells. We assembled a large biomolecular interaction knowledge base that can be leveraged to predict T cell phenotypes and guide CAR design. We also found that incorporating auxiliary search terms, such as biological process ontology terms, yields more relevant interaction papers than searching by protein names alone. This conclusion is based on the current machine readers and algorithms used in PubMed. Additionally, the generation from LLMs might include unexpected output, introducing failures in the extraction pipeline and increasing false negative rates. This limitation could be mitigated by using grammar-constrained decoding. Future work should also focus on evaluating the correctness of extracted triples and enriching the knowledge graph through the integration of additional data sources.

## ACKNOWLEDGEMENTS

This research was supported by the NSF EAGER award CCF-2324742, the NSF CAREER award CCF-2442884, and the University of Pittsburgh Center for Research Computing RRID:SCR_022735, through the resources provided. Specifically, this work used the H2P cluster, which is supported by NSF award number OAC-2117681.

## Appendix

### Queries

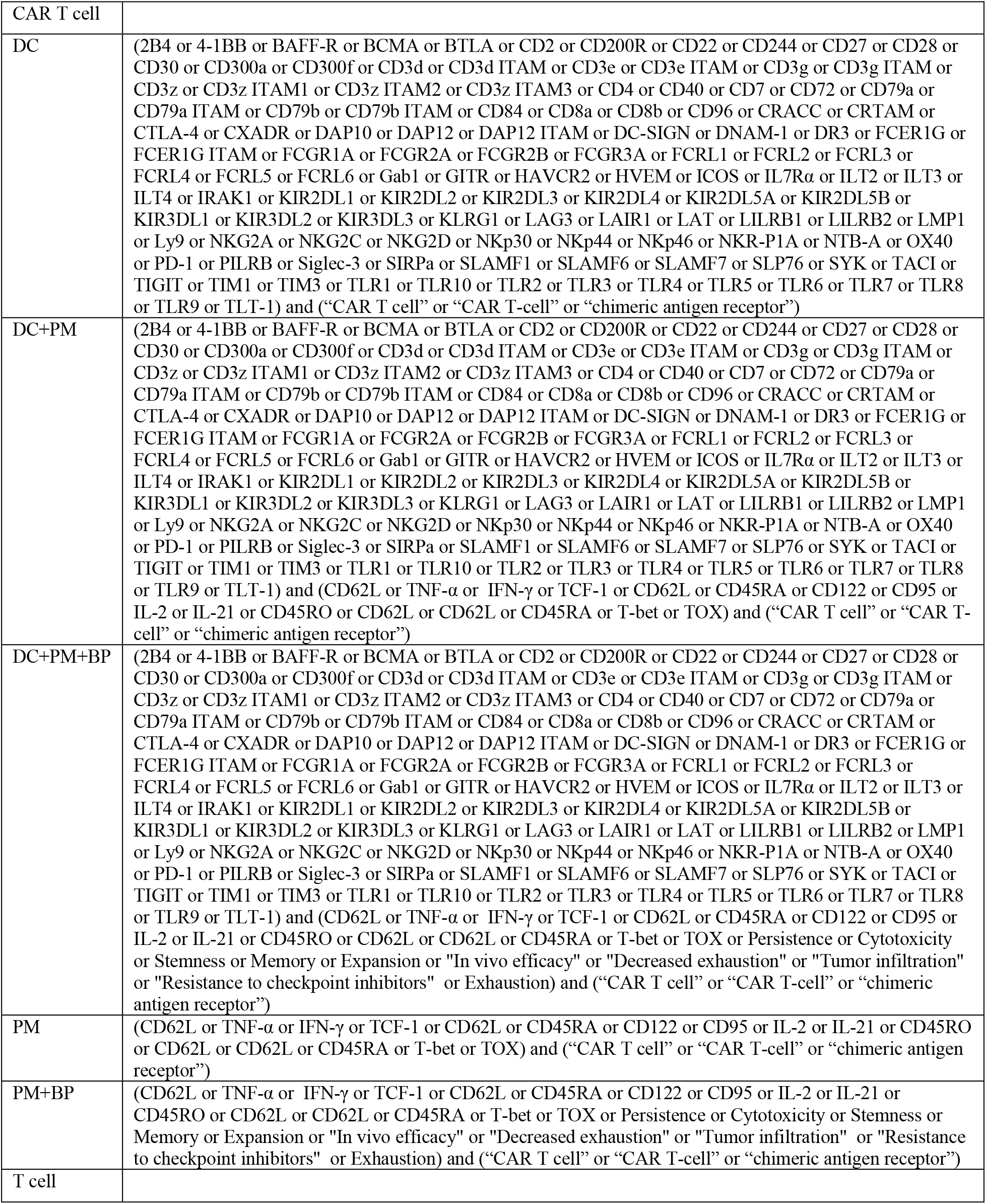
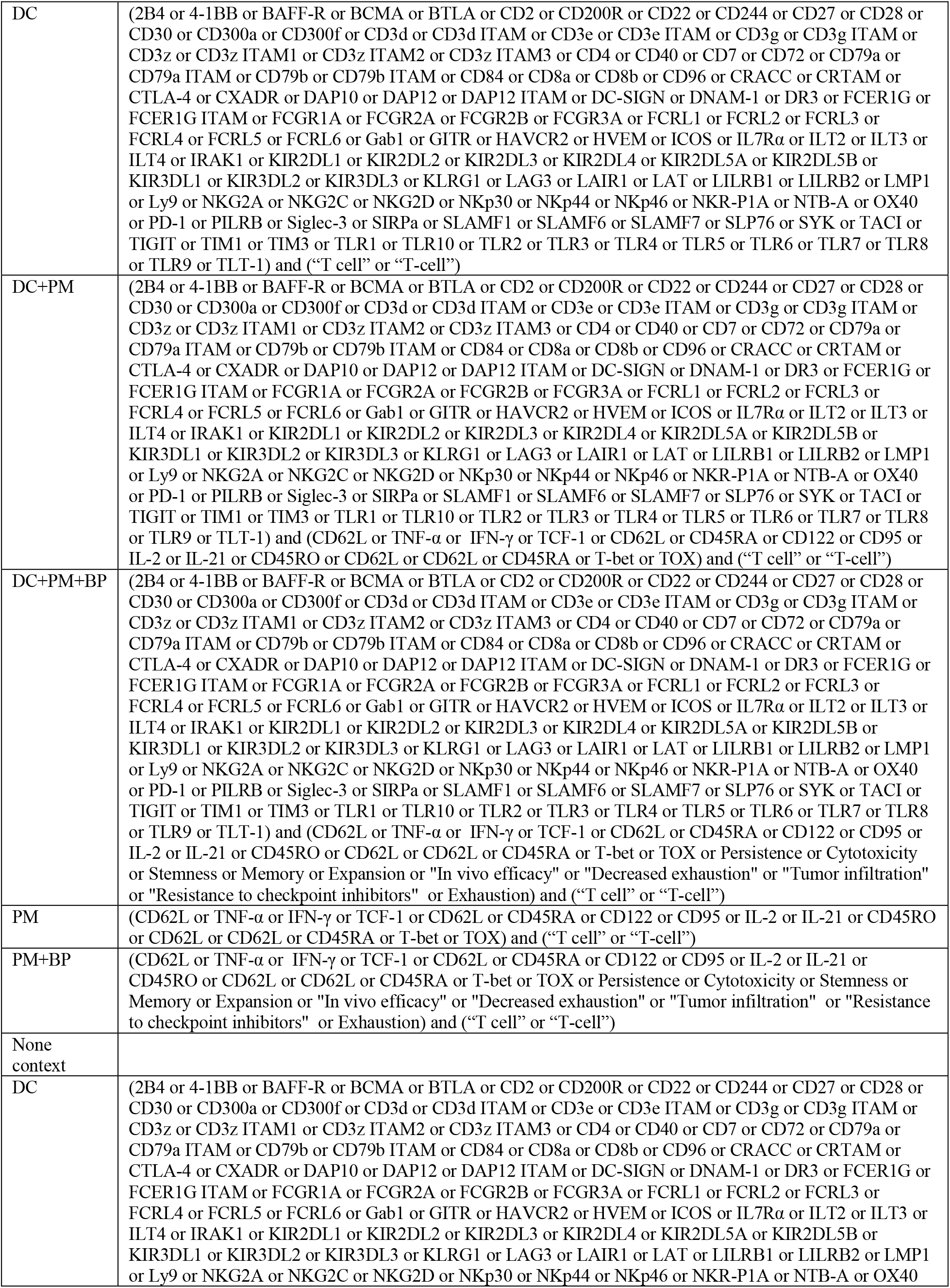
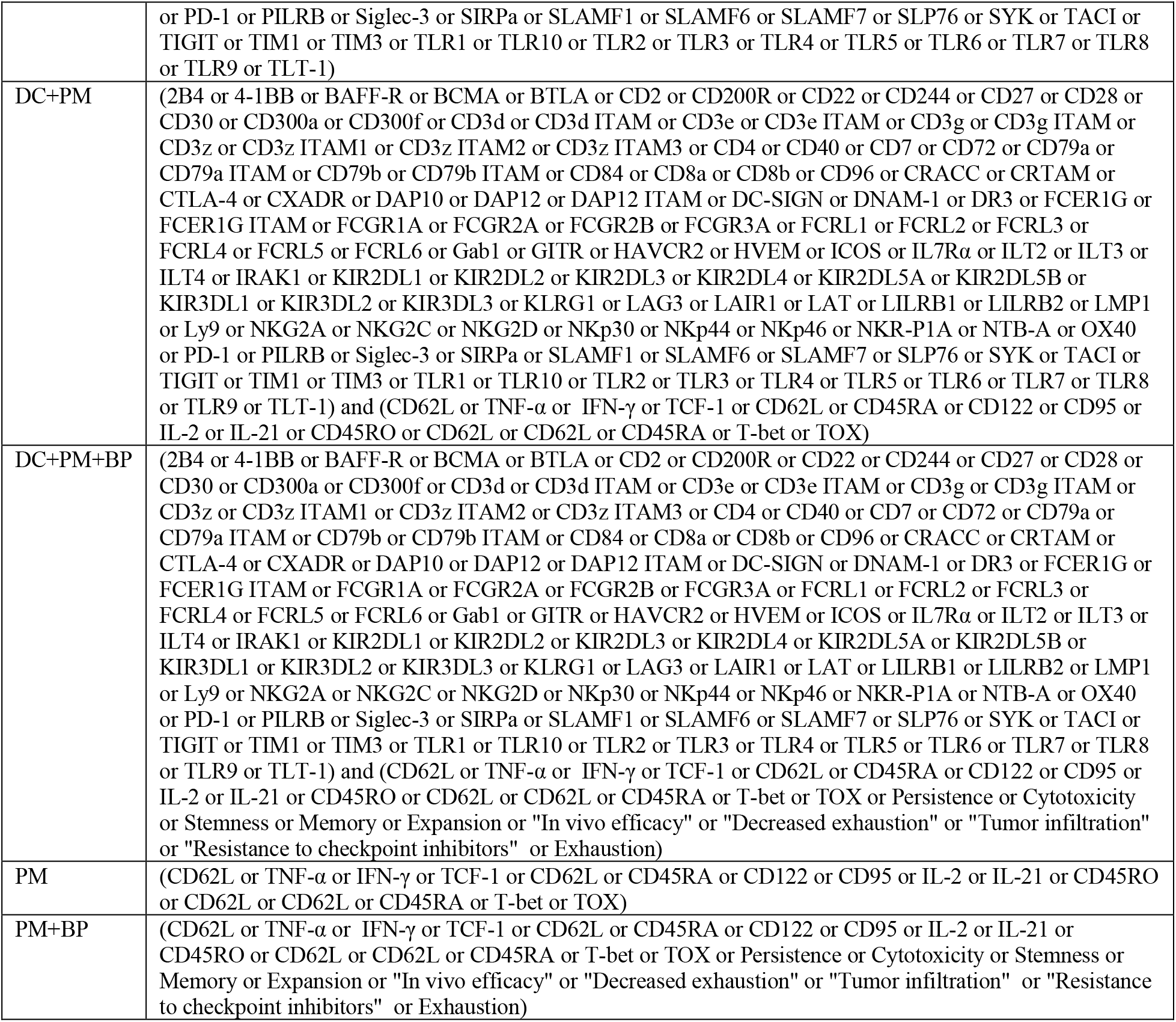

## REFERENCES

[1] D. W. Lee et al., “T cells expressing CD19 chimeric antigen receptors for acute lymphoblastic leukaemia in children and young adults: a phase 1 dose-escalation trial,” The Lancet, vol. 385, no. 9967, pp. 517–528, 2015.

[2] C. Zhang, J. Liu, J. F. Zhong, and X. Zhang, “Engineering car-t cells,” Biomarker research, vol. 5, pp. 1–6, 2017.

[3] K. S. Gordon et al., “Screening for CD19-specific chimaeric antigen receptors with enhanced signalling via a barcoded library of intracellular domains,” Nature biomedical engineering, vol. 6, no. 7, pp. 855–866, 2022.

[4] D. B. Goodman et al., “Pooled screening of CAR T cells identifies diverse immune signaling domains for next-generation immunotherapies,” Science translational medicine, vol. 14, no. 670, p. eabm1463, 2022.

[5] K. G. Daniels et al., “Decoding CAR T cell phenotype using combinatorial signaling motif libraries and machine learning,” (in eng), Science, vol. 378, no. 6625, pp. 1194–1200, Dec 16 2022, doi: 10.1126/science.abq0225.

[6] B. Landesman, “PubMed central,” Reference Reviews, vol. 19, no. 3, pp. 37–38, 2005.

[7] N. Fiorini et al., “PubMed Labs: an experimental system for improving biomedical literature search,” Database, vol. 2018, p. bay094, 2018.

[8] E. Holtzapple, B. Cochran, and N. Miskov-Zivanov, “Context-aware query design combines knowledge and data for efficient r eading and reasoning,” D. Demner-Fushman, K. B. Cohen, S. Ananiadou, and J. Tsujii, Eds.: Association for Computational Linguistics, 2021/6//, pp. 238–246, doi: 10.18653/v1/2021.bionlp-1.26. [Online]. Available: https://aclanthology.org/2021.bionlp-1.26

[9] A. R. Rivas, E. L. Iglesias, and L. Borrajo, “Study of query expansion techniques and their application in the biomedical information retrieval,” The Scientific World Journal, vol. 2014, no. 1, p. 132158, 2014.

[10] D. Tang, T. Y. C. Tam, H. Luo, C. A. Telmer, and N. Miskov-Zivanov, “An open-set semi-supervised multi-task learning framework for context classification in biomedical texts,” Journal of Biomedical Informatics, p. 104886, 2025.

[11] M. A. Valenzuela-Escarcega et al., “Large-scale automated reading with Reach discovers new cancer driving mechanisms,” in Proceedings of the Sixth BioCreative Challenge Evaluation Workshop, 2017, pp. 201–203.

[12] B. M. Gyori, J. A. Bachman, K. Subramanian, J. L. Muhlich, L. Galescu, and P. K. Sorger, “From word models to executable models of signaling networks using automated assembly,” Molecular systems biology, vol. 13, no. 11, p. 954, 2017.

[13] J. A. Bachman, B. M. Gyori, and P. K. Sorger, “Automated assembly of molecular mechanisms at scale from text mining and curated databases,” Molecular Systems Biology, vol. 19, no. 5, p. e11325, 2023, doi: 10.15252/msb.202211325.

[14] A. Dubey et al., “The llama 3 herd of models,” arXiv preprint 2407.21783, 2024.

[15] E. Holtzapple et al., “The BioRECIPE Knowledge Representation Format,” bioRxiv, p. 2024.02.12.579694, 2024.

[16] H. Luo et al., “VIOLIN: A modular framework for scalable reconciliation of heterogeneous interaction graphs,” bioRxiv, p. 2024.07.21.604448, 2026, doi: 10.1101/2024.07.21.604448.

[17] E. Holtzapple, C. A. Telmer, and N. Miskov-Zivanov, “FLUTE: Fast and reliable knowledge retrieval from biomedical literature,” Database, vol. 2020, p. baaa056, 2020.

[18] M. Kuhn et al., “STITCH 4: integration of protein–chemical interactions with user data,” (in en), Nucleic Acids Research, vol. 42, no. D1, pp. D401–D407, 2014/01// 2014, doi: 10.1093/nar/gkt1207.

[19] D. Szklarczyk et al., “The STRING database in 2023: protein–protein association networks and functional enrichment analyses for any sequenced genome of interest,” Nucleic acids research, vol. 51, no. D1, pp. D638–D646, 2023.

[20] M. Ashburner et al., “Gene ontology: tool for the unification of biology,” Nature genetics, vol. 25, no. 1, pp. 25–29, 2000.

[21] A. Hagberg and D. Conway, “Networkx: Network analysis with python,” URL: https://networkx.github.io, 2020.

